# Tracking molar wear in captive baboons: sex and age effects using a modified Scott scoring system

**DOI:** 10.1101/2025.06.12.659345

**Authors:** Kristin L. Krueger, Ian Towle, Gregory J. Matthews, Ana Álvarez Fernández, Leslea J. Hlusko

## Abstract

**Objectives:** This study evaluates molar wear progression in a captive baboon population under controlled dietary and environmental conditions. By comparing the dentin exposure ratio (DER) with a newly developed quadrant-based modification of Scott’s dental wear scoring system (Krueger-Scott method), we evaluate how wear patterns vary by age, sex, and occlusal region.

**Materials and Methods:** Mandibular second molars (M_2_) were assessed at two timepoints, during life and postmortem, in 201 captive baboons from the Southwest National Primate Research Center. Krueger-Scott and DER data were collected from 3D intraoral scans processed in MEDIT Link software. The Krueger-Scott scores assigned ordinal scores (1-10) to four equal quadrants of each M_2_ based on enamel facet development and dentin exposure. Statistical analyses tested relationships between wear progression, quadrant location, sex, and age.

**Results:** Krueger-Scott scores and DER values increased significantly between timepoints, indicating wear progression. However, Krueger-Scott scores revealed strong functional patterning: buccal and lingual cusps showed high within-group correlations and weaker cross-group correlations. Males showed significantly higher wear than females, despite being younger on average. The relationship between age and wear progression differed by sex.

**Discussion:** The Krueger-Scott method provided a more anatomically informative and efficient approach to tracking occlusal wear than DER. It captured regional wear variation and functional asymmetries that DER could not detect. Even under controlled conditions, sex-based differences in wear emerged, likely reflecting behavioral, morphological, or enamel structural variation. These findings offer a comparative baseline and demonstrate the utility of quadrant-level scoring for interpreting wear in extant and extinct taxa.

## Introduction

Teeth are powerful tools for reconstructing the biology, behavior, and ecology of living and extinct primates. While dental development is largely genetically controlled, post-eruptive wear is a complex interplay of intrinsic and extrinsic factors, including dietary composition, age, environment, occlusal morphology, and masticatory behavior (Lucas, 2004; Bartlett, 2005; Ungar, 2015). As teeth wear, they lose enamel, expose underlying dentin, and change morphology, affecting both chewing efficiency and long-term oral function. In primates, these wear processes are not merely passive, but are often adaptive responses to functional and ecological demands (e.g., King et al., 2005; Galbany et al., 2014; Pampush et al., 2016, 2018; Avià et al., 2022; Teaford et al., 2021, 2022; Pampush, Morse, & Kay, 2024).

Despite its importance, dental wear remains challenging to quantify and interpret. Most primate dental wear studies rely on cross-sectional data, which offer only a single snapshot of wear over time (e.g., Kay and Cant, 1988; Krueger et al., 2008; Elgart, 2010; Morse et al., 2013; Spradley, Glander, & Kay, 2016; Pampush et al., 2016, 2018; Galbany et al., 2011, 2020; Selig, Kupczik, & Silcox, 2021; Fiorenza et al., 2022; Guatelli-Steinberg et al., 2022; Harty et al., 2022; Pampush, Morse, & Kay, 2024; Selig et al., 2025). Longitudinal studies of wild primates are rare and often confounded by ecological variability, making it difficult to isolate the factors driving wear (e.g., Frohelich, Thorington, & Otis, 1981; Teaford and Glander, 1996; Phillips-Conroy, Bergman, & Jolly, 2001; Dennis et al., 2004; King et al., 2005; Wright et al., 2008; Galbany et al., 2011; Cuozzo et al., 2014). Controlled, longitudinal datasets, especially those involving well-documented individuals, are therefore critical for untangling intrinsic from extrinsic influences of wear (e.g., Teaford et al., 2021, 2022; Towle et al., 2024).

This study uses a unique two-timepoint dataset from a large sample of captive baboons (*Papio* sp.) housed under stable environmental and dietary conditions. To evaluate wear progression, we used two different approaches: 1). dentin exposure ratio (DER), a standard, continuous metric widely used in primate dental wear research (e.g., Kay & Cant, 1988; Phillips-Conroy et al., 2001; Elgart, 2010; Galbany et al., 2011, 2014, 2020; Morse et al., 2013; Pampush et al., 2016, 2018; Spradley et al., 2016; Selig et al., 2021, 2025) and 2) a novel quadrant-based modification of Scott’s (1979) ordinal wear scoring system, here called the Krueger-Scott method.

DER quantifies the proportion of the occlusal tooth crown covered by exposed dentin and is useful for capturing total wear. However, it treats the occlusal surface as a single unit and is less effective at identifying early enamel loss or regional variation. Moreover, comparing DER across species is difficult due to variation in cusp height, occlusal morphology, and enamel thickness (e.g., Martin, Olejniczak, & Maas, 2003; Galbany et al. 2020; Morse, Pampush, & Kay, 2023).

The Krueger-Scott method addresses these limitations by scoring four quadrants of the molar independently on a 1-10 ordinal scale, allowing for detection of fine-scale variation in enamel loss and dentin exposure (Figure 1). Adapted from Shykoluk and Lovell (2010), this method is cost-effective, replicable, and usable on both dental casts and skeletal material. It captures both enamel and dentin wear, aligns with functional zones of the tooth, and facilitates meaningful comparisons across bilophodont taxa. Research in occlusal topography confirms that cusps do not wear evenly and differential wear can reflect underlying biomechanical function (Ungar and M’Kirera, 2003; Glowacka et al., 2016, Towle et al., 2021; Avià et al., 2022). Indeed, quadrant-level scoring can reveal subtle asymmetries and functional shifts in occlusal loading that whole-tooth metrics like DER would overlook.

**Figure 1.**
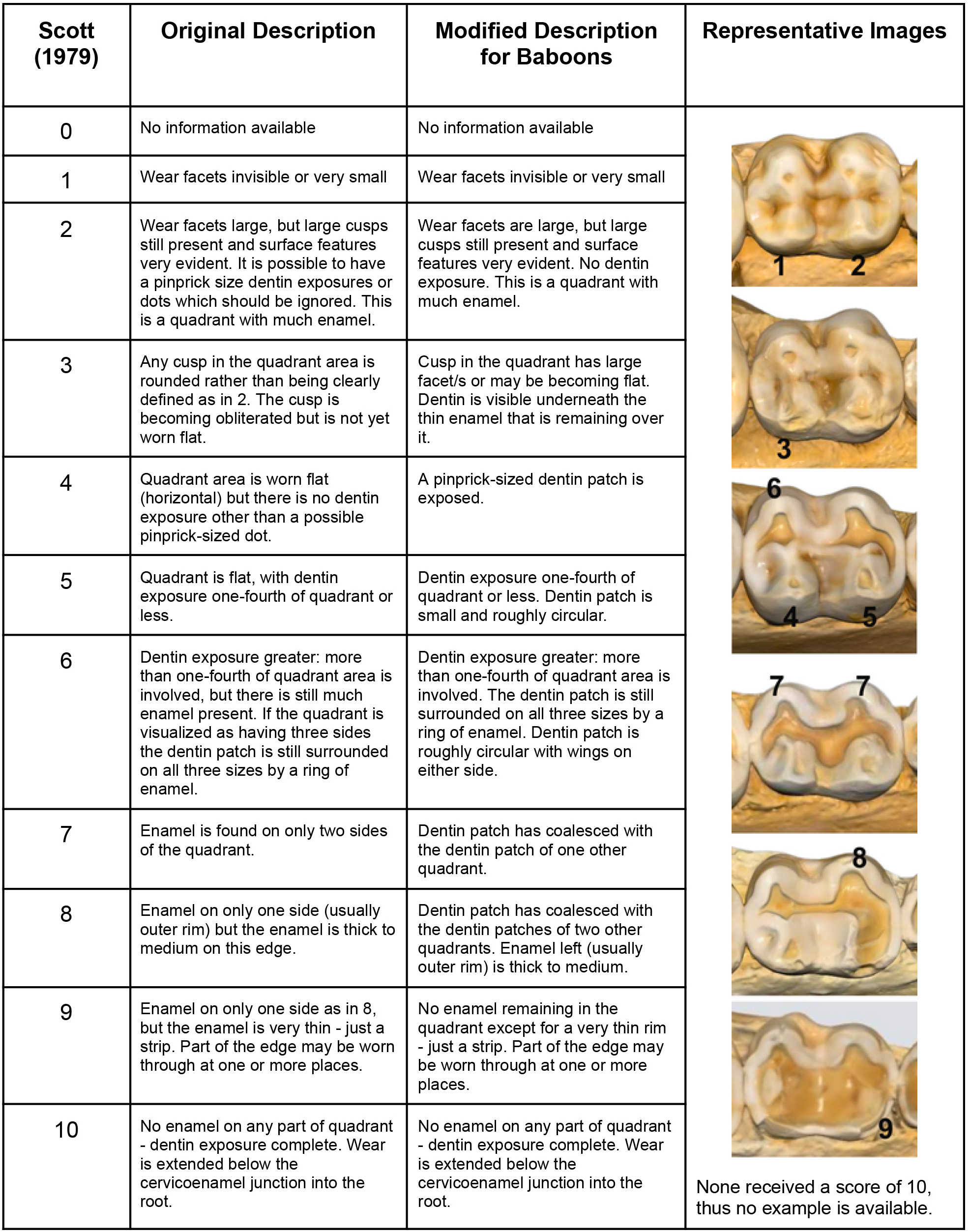
Original Scott score descriptions (taken from Scott, 1979), modified descriptions for use in baboons, and representative images of each modified wear score.

Research on baboon dental wear has yielded important insights into their dietary behavior, ecological conditions, and life history patterns (e.g., Bramblett, 1969; Phillips-Conroy et al., 2001; Nystrom et al., 2004; Galbany et al., 2011; Avià et al., 2022). Most of this work, however, has focused on wild populations and relied heavily on cross-sectional data and using percent dentin exposure (PDE or DER x 100) as the primary wear metric. These studies have appropriately emphasized diet and environmental abrasives as the primary drivers of wear variation. For example, Phillips-Conroy et al. (2001) compared PDE in wild baboons from Mikumi National Park, Tanzania and Awash National Park, Ethiopia, finding population-level differences in wear rate. They attributed these differences to variation in dietary abrasiveness, concluding that both age and environmental context shaped molar wear progression.

Galbany et al. (2011) examined PDE in Amboseli baboons and found that age and the ingestion of grit-covered grass corms were the main contributors to wear, based on detailed behavioral and ecological data. A follow-up study comparing mandrills and baboons revealed faster wear rates in mandrills, attributed to higher quartz content in their environmental sediments (Galbany et al., 2014). While sex differences in wear were not observed in these studies, dietary and sedimentary variation clearly influenced occlusal wear patterns. Similarly, microwear analyses of baboons from Awash National Park showed patterns consistent with tough, abrasive diets, although no significant effects of age, sex, or troop were found (Nystrom et al., 2004).

Despite their contributions, these studies share important limitations. Most are cross-sectional, rely on PDE or DER as whole-tooth metrics, and are conducted in variable ecological contexts. As noted by Galbany et al. (2020), PDE may miss early stages of wear, can be difficult to interpret across species with different occlusal morphologies, and is most effective when applied to individuals of known age. These limitations underscore the need for controlled, longitudinal datasets using methods that can detect intra-tooth variation.

Our study addresses these gaps by combining DER and a novel quadrant-based scoring system to analyze molar wear progression in captive baboons raised under uniform dietary and environmental conditions. This unique dataset allows us to assess how wear progresses over time within individuals, evaluate the effects of age and sex, and explore occlusal regionality in wear patterns.

Here, we test four hypotheses:

1. Wear increases over time. We expect both Krueger-Scott scores and DER values to increase between casting and death.
2. Quadrants are functionally structured. We expect quadrant-level Krueger-Scott scores to be positively correlated, especially within buccal and lingual cusp groups.
3. There are no sex differences in wear patterns. We anticipate that males and females show similar wear patterns due to uniform diets.
4. The relationship between age and wear progression does not differ by sex. We expect wear accumulates at the same rate across the lifespan for both males and females.

This study provides a new framework for understanding molar wear under controlled conditions. By integrating whole-tooth and quadrant-level approaches, we aim to clarify the drivers of wear progression, offer a reference point for interpreting wear in wild or fossil samples, and advance the toolkit available for primate dental research.

## Materials and Methods

This study analyzed mandibular second molars (M_2_) from 201 captive baboons, derived from a larger, captive population housed at the Southwest National Primate Research Center (SNPRC) in San Antonio, Texas, USA. Established in the early 1970s, the SNPRC colony spans nine generations and serves as a pedigreed population supporting quantitative genetic analysis of a variety of phenotypes, including metabolic (Higgins et al., 2010; Mahaney et al., 2018; Lin et al., 2024), behavioral (Johnson et al., 2015), life history (Martin et al., 2002), skeletal (Hansen et al., 2009; Havill et al., 2010), and dental (Koh et al., 2010; Hlusko et al., 2016) traits. All animals used here were born and reared at SNPRC, housed in large, outdoor social enclosures, and fed a consistent diet of commercial monkey chow and water (Hlusko, Mahaney, & Weiss, 2002).

Baboon taxonomy remains complex due to widespread hybridization in the wild (Kopp et al., 2023). While modern baboons are typically classified as six subspecies within *Papio hamadryas* (Jolly, 1993), or more commonly today, as six separate species (*P. papio*, *P. hamadryas*, *P. anubis*, *P. cynocephalus*, *P. kindae*, and *P. ursinus*; Elton and Dunn 2020), historical classifications, including those applied to the SNPRC colony, often treated them as subspecies of *Papio hamadryas*. Founding individuals of the SNPRC colony originated from the *hamadryas*, *anubis*, and *cynocephalus* taxa, and the colony has been described using this taxonomy in earlier work (e.g., Hlusko et al., 2016).

Molar wear for each individual was evaluated at two time-points: 1) during life, from high-resolution dental casts of adult dentitions, and 2) postmortem, from skeletal remains. Life-stage casts were derived from dental molds collected according to the *Guide for the Care and Use of Laboratory* Animals (National Research Council, 1996) and approved by both the SNPRC and University of Illinois Animal Care and Use Committees (see Hlusko et al., 2002). Plaster replicas of these molds are housed at the Centro Nacional de Investigación sobre la Evolución Humana (CENIEH) in Burgos, Spain (for details, see: Hlusko et al., 2002; Hlusko and Mahaney, 2003; Hlusko, Maas, and Mahaney, 2004). Postmortem specimens were collected after necropsy by Dr. James Cheverud and curated in the Department of Biology at Loyola University Chicago in Chicago, USA. They are now housed at the University of Illinois, Urbana-Champaign, under the care of Dr. Charles Roseman.

To ensure longitudinal consistency, all individuals included in this study had dental casts collected within five years prior to death, with intervals ranging from 0.05 to 4.96 years (Krueger et al., 2025). The left M_2_ was prioritized; when absent or damaged, the right M_2_ was used following standard anthropological protocol (Buikstra and Ubelaker, 1994). Only one tooth per individual was scored. While bilateral comparisons may offer insight into chewing-side preference, we opted for a single-tooth approach to maximize sample size and avoid overrepresenting individuals with more complete remains.

Occlusal surfaces of both timepoints were digitized using a MEDIT i700 intraoral scanner, which generates high-resolution 3D models in STL format, suitable for surface area measurements. DER was calculated using the area measurement tool within the MEDIT Link Design application within the MEDIT Link software v 3.4.2 (Medit, Seoul, Republic of Korea). For each scan, observers manually delineated the exposed dentin area and total occlusal surface (Figure 2). DER was defined as the ratio of exposed dentin area to total occlusal area: DER = (dentin area / total occlusal area). This measurement was collected for both timepoints by two observers (AAF and BDE) and using a protocol developed by KLK and adapted from prior studies (e.g., Galbany et al., 2011; Spradley et al., 2016).

**Figure 2.**
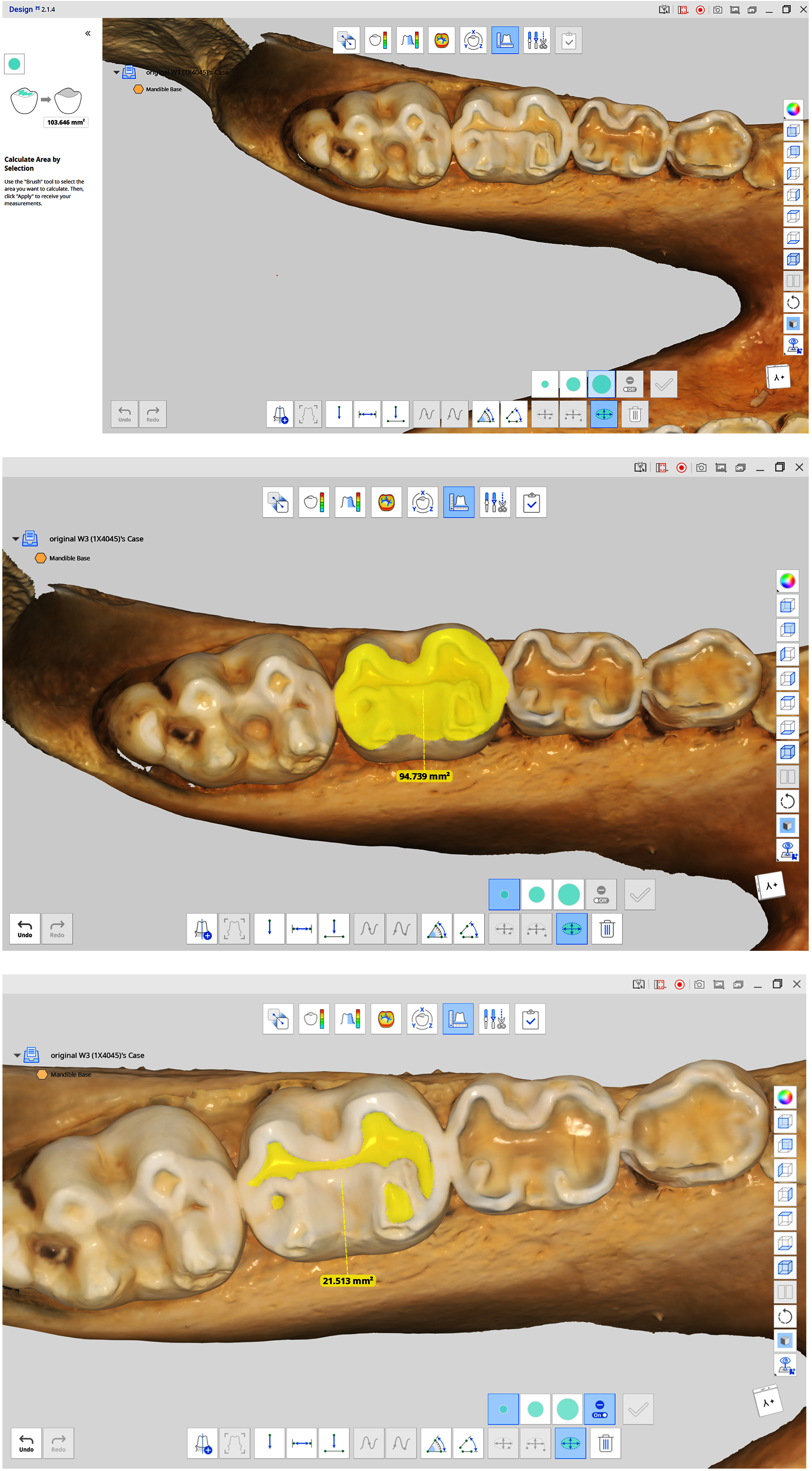
Representative images of DER measurements using MEDIT measurement tools. Top image shows the tooth prior to measurement, middle image shows occlusal surface area measurement, and bottom image shows dentin area measurement.

Traditional DER methods typically rely on 2D techniques, such as hand-drawn occlusal outlines or photographic images (e.g., Spradley et al., 2016; Galbany et al., 2020), to calculate the proportion of exposed dentin relative to the crown surface visible in occlusion. However, this introduces a common limitation found in dental microwear studies: two-dimensional projections are being used to represent inherently three-dimensional structures (Ungar et al., 2008; Krueger, 2015). In contrast, our 3D approach allowed observers to zoom, rotate, and examine the molar surface interactively (Figure 2). This reduced project-related distortion and enabled measurement of dentin exposure relative to the full occlusal landscape rather than the crown silhouette. While our method focused on surface area, future work could incorporate 3D volumetric assessments to further refine these measures (Towle et al., 2024). DER scores were collected twice for each individual. This process was divided between two technicians over two months and is estimated to have required just under 100 person-hours in total.

We also evaluated occlusal wear using a modified version of Scott’s (1979) ordinal wear scoring system, adapted for bilophodont cercopithecoid molars (Figure 1). Each molar was divided into four equal quadrants: mesiobuccal (MB), distobuccal (DB), mesiolingual (ML), and distolingual (DL), using the waist of the talonid basin and the nadirs of the transverse lophs as reference points. While quadrant names align with standard cusp terminology, the entire quadrant (not only the cusp) was scored.

Wear scores ranged from 1 (unworn) to 10 (near-complete enamel loss with substantial dentin exposure). This modified approach, developed for baboon morphology, was based on the quadrant-level protocol described by Shykoluk and Lovell (2010), rather than the simplified systems used by Avià et al. (2022) and Conrad (2023). All Krueger-Scott scores were assigned by a single observer (KLK) over a two-week period to maintain internal consistency across the dataset.

To test Hypothesis 1, for Krueger-Scott scores, two-sample dependent t-tests were used to compare the mean wear scores for each quadrant at casting and at death. This was done to determine if the mean wear scores at casting and at death were different at individual quadrants. A Bonferroni correction was used to control the family-wise error rate (FWER) across the 5 tests compared here. For DER, since we had two observers, a mixed-effects ANOVA was used to compare the mean wear scores at casting and at death. We included a random effect for the observer.

To test Hypothesis 2, Spearman’s correlation coefficients were computed for all pairs of quadrants and timepoints (i.e., cast and skeletal). Spearman’s method was used to avoid the assumption of linearity and instead measure the strength and direction of monotonic relationships. Comparing all possible pairs of quadrants and times to all others results in 28 tests, and a Bonferroni correction was used to control the FWER at level 0.05.

To test Hypothesis 3, for Krueger-Scott scores, we performed Hotelling’s two-sample T^2^ test to assess differences in the mean wear vectors for males versus females at casting and death. As we rejected the null hypothesis, post-hoc tests (i.e., Welch’s t-tests) were used to conduct pairwise comparisons for each of the eight individual quadrant-time combinations (e.g., mesiobuccal cast score, mesiobuccal skeletal score, etc.) to determine the location of significant mean differences between sexes. A Bonferroni correction was used to control the family-wise error rate at level 0.05 across the eight tests performed here. For DER, since we had two observers, a mixed-effects regression model was used with sex as the fixed effect and observer and baboon as the random effect.

For Hypothesis 4, two mixed-effects regression models were fit, a full and reduced model, for each measure of wear. For the Krueger-Scott wear measure, both the full and reduced models had the same random effect, which were random intercepts for baboons to account for individual-level variation. The full model had fixed effects for sex, age, age^2^, age^3^ and interactions between sex and age, age^2^, age^3^. The reduced model included only age, age^2^, age^3^. For the DER wear measure, all the model parameters were the same, except both the full and reduced models had an additional random intercept for the observer.

Likelihood ratio testing was used to compare the full model to a reduced model. In this analysis, we included three timepoints for each baboon even though wear scores were only universally measured at two timepoints for each individual. This third timepoint comes from assuming that the Krueger-Scott wear score for each individual quadrant at eruption is one, and the DER value for each individual baboon is zero, and we assume that eruption for the M_2_ occurs at 42 months, which is the average time for baboons. (PhillipsLConroy and Jolly, 1988).

For visualization purposes, we used isotonic regression (Robertson, Wright, & Dykstra, 1988) to estimate wear curves for males and females separately. Isotonic regression requires the functional form of the relationship to be monotonically non-decreasing, fitting well with wear scores that cannot decrease with age. We also assessed the intraclass correlation (ICC) for each Krueger-Scott wear quadrant score, Krueger-Scott sum score, DER by observer, and DER by individual baboon using data from three timepoints: eruption, casting, and death (Bartko, 1966; Müller and Büttner, 1994). The ICC is a statistical measure that quantifies the similarity between individuals within a group. The ICC was calculated as the proportion of variance attributable to individual baboon ID, defined as the variance due to the ID divided by the sum of ID and residual variance. It was also calculated for both DER observers (i.e., interobserver error). All statistical analyses were performed in R, using add-on packages LME4, Iso, Hotelling, ggplot2, and tidyverse (version 4.2.2; R Core Team, 2024; Bates et al., 2015; Curran and Hersh, 2021; Turner, 2023; Wickham, 2016).

## Results

Data for this study are available open access through the Dryad data repository (<u>Krueger</u> <u>et al., 2025</u>). Descriptive statistics are presented in Tables 1, 2a, and 2b. Across all individuals, Krueger-Scott scores (summed and by quadrant) and DER increased between the life (cast) and death (skeletal) timepoints, consistent with wear accumulation through time. Mean summed and quadrant-level Krueger-Scott scores were significantly higher at death than at casting across all analyses (Table 2a). Paired-sample t-tests revealed statistically significant increases for the summed score and by quadrant (Table 3a). Bonferroni correction for multiple comparisons maintained the significance of all results. Similarly, average DER at casting was 0.118 (sd=0.107) and at death was 0.151 (SD = 0.126, Table 2b). An ANOVA indicated this increase was statistically significant (Table 3b). These findings confirm that Krueger-Scott scores and DER increased significantly from casting to death (Hypothesis 1).

Spearman correlation coefficients showed strong positive correlations between quadrant wear scores within both timepoints (Table 4). Correlations were especially high within functional cusp groups: the buccal quadrants (MB and DB) were strongly correlated at both casting and death, as were the lingual quadrants (ML and DL). In contrast, buccal-lingual quadrant correlations were consistently weaker. All pairwise comparisons between quadrant and timepoint combinations showed positive correlations, with varying strengths. After Bonferroni correction, all comparisons remained statistically significant. These results support Hypothesis 2, indicating that wear accumulates more synchronously within regions of similar occlusal function (buccal or lingual) than across them.

Both Krueger-Scott scores and DER were significantly different by sex. For Krueger-Scott scores, Hotelling’s T^2^ tests indicated significant sex-based differences in the mean vector of wear at both casting and death (Table 5a). Despite males being younger on average than females (Table 1), they showed significantly higher wear scores in multiple quadrant-timepoint combinations. Welch’s t-tests found significant differences between males and females for all lingual quadrants (Table 5b). However, none of the buccal quadrants were found to be significant. The mixed effects regression model found significant differences in DER between the sexes at casting and at death (Table 6). Indeed, average DER at casting was 0.152 for males and 0.106 for females and at death, 0.206 for males and 0.132 for females (Table 2b). These findings refute Hypothesis 3 and suggest that wear progression is shaped by both age and sex-specific patterns.

To assess the influence of age, sex and wear progression, we used a mixed-effects regression model. A likelihood ratio test was used to test whether sex was a significant predictor in a model controlling for age (including a quadratic and cubic term). Results showed a significant difference in wear between males and females after controlling for age. This was consistent for all four Krueger-Scott quadrants, their sum, and DER (Tables 7a and 7b). The fitted isotonic regression curves (Figures 3 and 4) further illustrated that wear was consistently greater in males than in females in all cases. These results refute Hypothesis 4 and indicate that wear progression is not only age-related but differs by sex in its trajectory.

**Figure 3.**
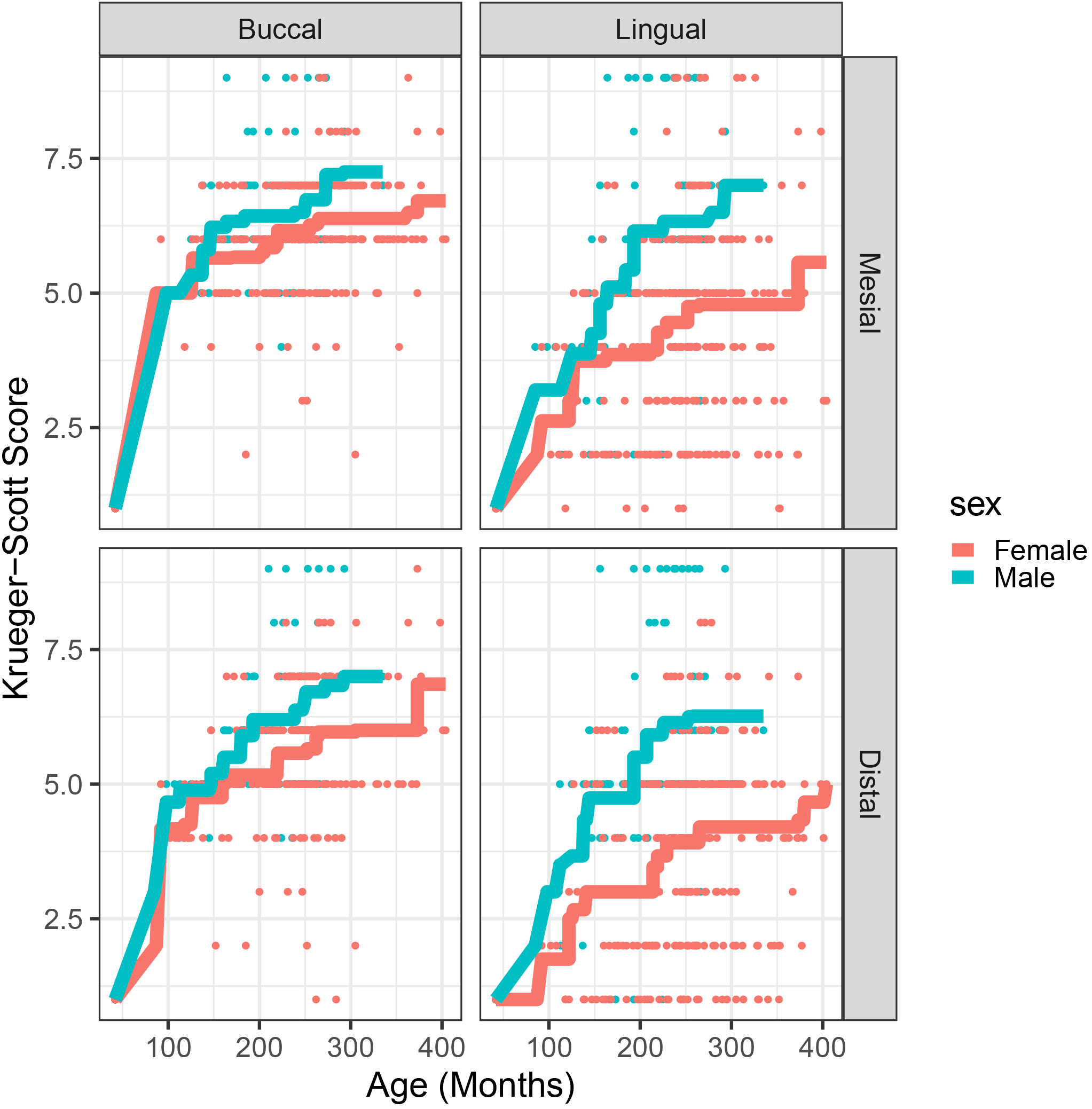
Isotonic regression of individual quadrants by sex. Krueger-Scott scores are on the y-axis, age (in months) on the x-axis.

**Figure 4.**
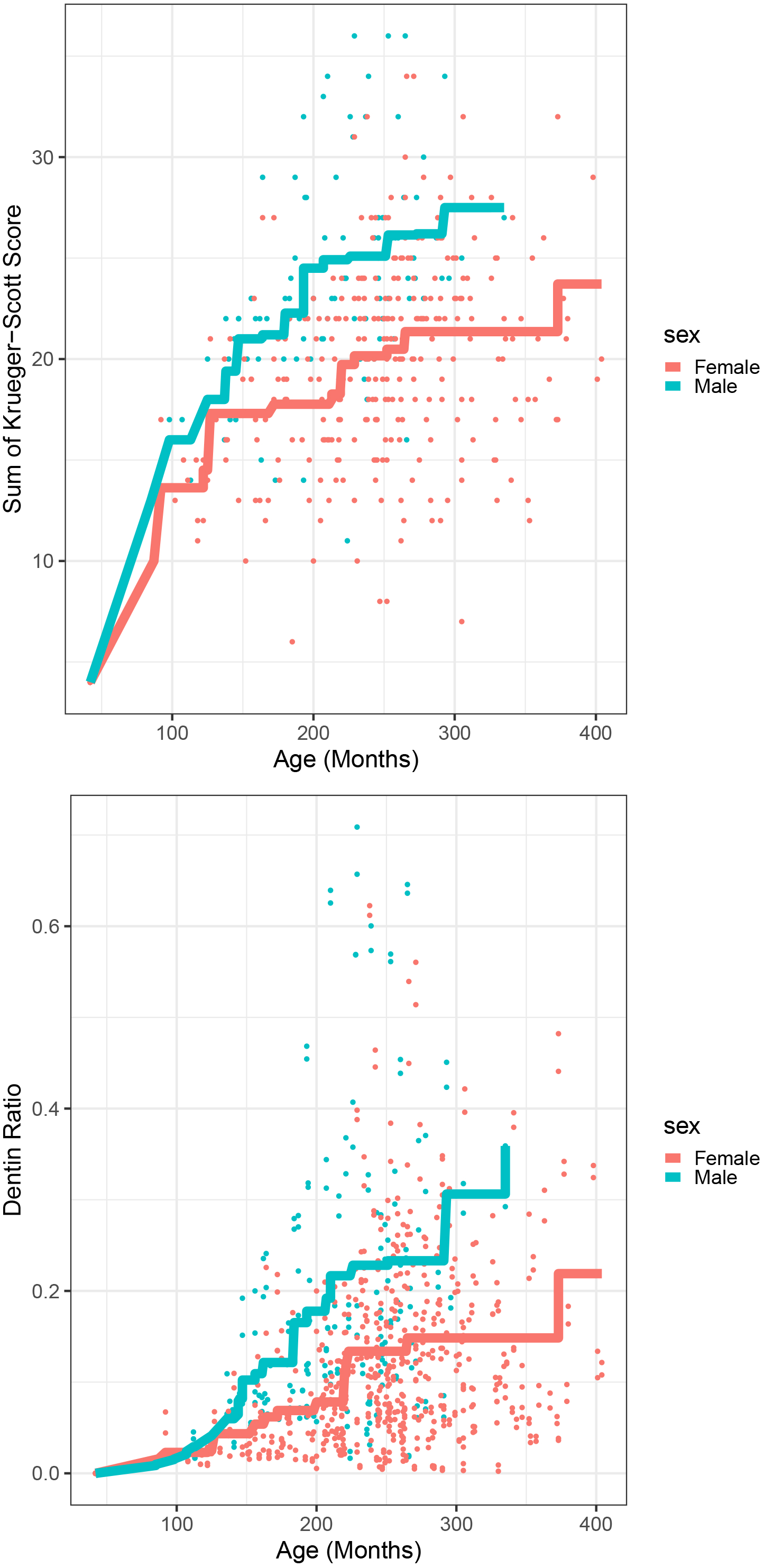
Isotonic regression of summed Krueger-Scott scores and DER values by sex. Wear measure is on the y-axis, age (in months) on the x-axis.

Finally, we assessed intraclass correlation (ICC) for each Krueger-Scott quadrant, Krueger- Scott sum, DER by observer, and DER by individual baboon (Tables 7a and 7b), using data from three timepoints: eruption, casting, and death (Bartko, 1966; Müller and Büttner, 1994). All four Krueger-Scott quadrants, as well as the summed score, showed similar ICC values of approximately 0.3, indicating that about 30% of the total variance in wear scores is explained by differences among individuals (Table 7a). For DER, the ICC by observer was approximately 0.002 while the ICC by individual ID was 0.546, suggesting that measurement differences between observers accounted for only 0.2% of the variance, whereas individual baboons accounted for 55% (Table 7b). While it may be tempting to directly compare ICC values across methods, these values are derived from different models and are therefore not directly comparable.

## Discussion

Deciphering dental wear is crucial for understanding the evolutionary and environmental adaptations of many organisms. Dental wear results from a complex interplay of factors, including age, diet, ecological conditions, mechanical and physical properties of enamel, and potentially other unknown influences. Minimizing the variability of these factors can help disentangle their specific effects. In this study, we used a large sample of captive baboons raised under uniform environmental conditions and diets, thereby reducing extrinsic sources of variation. We quantified their dental wear using a new quadrant-based version of Scott’s ordinal scoring system (the Krueger-Scott method) and DER to assess how wear progresses across time, age, and sex.

### Krueger-Scott scores and DER track wear progression through time

Both quadrant-level Krueger-Scott scores and DER showed statistically significant increases in molar wear between the life (cast) and death (skeletal) timepoints (Tables 3a and 3b). This confirms that wear progression can be reliably detected using either method, even across relatively short intervals. These increases support the utility of the Krueger-Scott method for tracking regional changes in occlusal morphology and validate the use of DER as a global wear metric. The consistency between methods also underscores the reliability of dental wear as a time-sensitive signal, even in a controlled sample where diet and environmental conditions were held constant.

DER quantifies the overall extent of occlusal wear but does not capture where that wear occurs. The Krueger-Scott method, on the other hand, provides a finer anatomical resolution by isolating quadrant-level changes.

### Occlusal wear patterns are functionally structured

Molar wear did not accumulate evenly across the occlusal surface. This study revealed substantial and patterned variation in wear between occlusal regions that over the lifespan, highlighting the dynamic and functionally complex nature of dental wear in primates. As expected, quadrant wear scores were positively correlated at both timepoints, likely due to their shared occlusal environment (Table 4). However, the strongest correlations were consistently found within buccal quadrants (mesiobuccal and distobuccal) and within lingual quadrants (mesiolingual and distolingual), suggesting that wear accumulates more synchronously within these functional groupings than across them.

Across all individuals, buccal cusps consistently showed higher wear scores than lingual cusps, regardless of age or sex. However, the greatest increase in quadrant scores over time occurred in the lingual cusps (Table 2a), suggesting that buccal cusps undergo more substantial wear earlier in life. This aligns with previous research showing that mandibular buccal cusps, due to their prominent role in initial food processing and occlusion with opposing maxillary molars, are subject to greater abrasive and compressive forces (Lucas, 2004). In baboons, the second molars erupt between three and four years of age (PhillipsLConroy and Jolly, 1988), and even among the youngest individuals in our study (7-10 years old), buccal quadrants already reached moderate scores (typically four or five; Figure 3). These patterns reflect well-documented phase-based wear trajectories in primates, shaped by enamel thickness, cusp architecture, and the biomechanics of the masticatory cycle (e.g., Molnar and Gantt, 1977; Teaford, 1982; Knight- Sadler and Fiorenza, 2017; Towle et al., 2023).

Lingual cusps, in contrast, showed a pronounced “catch-up” in wear between timepoints (Table 2a). This delayed but accelerated wear in lingual cusps observed later in life is unlikely to be linked to dietary changes, as all individuals were fed the same diet throughout their lives, and weaning typically occurs prior to M_2_ eruption (Lewis et al., 1988; Altmann, 2001). Rather, shifts in occlusal forces across the crown as dental wear progresses likely drive this change. This scenario is supported by human dental studies showing that the occlusal plane changes over time, with posterior teeth flattening and shifting wear toward lingual cusps as enamel is lost and occlusal morphology changes (Osborn, 1982; Kaifu, 1999). Our results align with this pattern and suggest a similar adaptive mechanism in baboons, whereby altered force distribution helps maintain masticatory efficiency despite cumulative tissue loss (Ungar and M’Kirera, 2003; Lucas, 2004; Galbany et al., 2011).

Yet, changes in occlusal plane and force vectors alone may only partially explain the greater variability observed in lingual cusp scores at both timepoints. While buccal wear tended to follow a more consistent trajectory, the Krueger-Scott scores for lingual quadrants showed more between-individual variation (Table 5b). One potential explanation is differential cusp fracture. Previous studies show that fractures occur more frequently on non-functional than functional cusps in a wide range of primates (Towle et al., 2021), and this pattern appears to hold true for this sample. Lingual cusp chipping may elevate wear scores in some individuals while leaving others unaffected, contributing to the increased variability observed in Krueger-Scott scores across lingual quadrants. This possibility highlights how micro-damage, even in the absence of dietary change, can influence macroscopic wear patterns.

These patterns also raise important questions for future research, particularly regarding the potential genetic correlations among cusp regions and their susceptibility to wear or fracture. Given prior evidence for heritable variation in cusp morphology in this same baboon population (Koh et al., 2010), further investigation could clarify the extent to which observed wear variation reflects biomechanical loading, genetic architecture, or both.

### Sex differences in wear despite a controlled diet

Despite a uniform diet and housing conditions, male baboons in our sample had greater dental wear than females. This finding held for Krueger-Scott quadrant scores and DER values (Tables 5a and 6). Surprisingly, males were also younger on average than females, yet still showed significantly more occlusal wear, indicating that they wore down their teeth more rapidly and to a greater extent. These findings align with observations in wild baboons in Kenya, where males also showed greater occlusal wear than females (Bramblett, 1969), although that study attributed the pattern to higher female mortality due to predation, a factor irrelevant to our captive sample. On the other hand, our results differ from other studies on wild baboons, which reported minimal (Avià et al. 2022) or no sex differences in occlusal wear (Galbany et al., 2014). Thus, specific factors within this captive sample may help explain the consistent sex differences found here.

One possible explanation is behavioral. Anecdotal reports from one of the co-authors (LJH), who collected the dental molds, noted that some baboons softened their monkey chow by dipping it in water before chewing. This behavior would reduce the mechanical force required during mastication, possibly reducing enamel wear. Although the specific individuals engaging in this behavior were not recorded, it is possible that it occurred more frequently in females which could contribute to sex differences in occlusal wear. However, without systematic behavioral data, this remains speculative.

Another behavioral factor may involve stress-induced bruxism. Captive primates, especially males, are known to display abnormal behaviors such as hair pulling, self-aggression, and teeth grinding under conditions of social stress or confinement (Bellanca and Crockett, 2002; Rommeck et al., 2009; Gettleman, 1981; Gottlieb, Capitanio, and McCowan, 2013, but see Brent, 2009). Although SNPRC baboons were housed in social groups with established social and dominance hierarchies (O’Connor et al., 2011), they lacked the ecological challenges of foraging or predator avoidance, potentially increasing the emphasis on social interactions (Brent, 2009). Studies showed that social crowding in baboons, whether chronic or acute, elevated salivary cortisol levels (Pearson, Reeder, and Judge, 2015), and it is possible that elevated stress in males contributed to increased wear through parafunctional activity such as bruxism.

In addition to behavioral influences, enamel structure may play a role. Quantitative genetic analyses in the SNPRC colony found that *relative* molar enamel thickness (enamel thickness scaled by a measure of tooth size) is sexually dimorphic, whereas *absolute* enamel thickness is not (Hlusko et al., 2004). Given that tooth size is also sexually dimorphic, with sex accounting for 28% of phenotypic variation in molar size (Hlusko, Sage, & Mahaney, 2011), this suggests that males have a greater volume of dentin beneath a proportionally thinner enamel layer. This histological configuration may leave male molars more vulnerable to wear once enamel is breached, potentially accelerating wear rates.

Physical differences related to colony management may also be relevant. In captive settings, male baboons frequently undergo blunting or removal of maxillary canines to reduce the risk of injury to conspecifics and handlers (Smith, 1971; Tomson, Schulte, and Bertsch, 1979; Schofield et al., 1991) and due to canine fracture (Facchini et al., 2018). In our sample, 48 of 51 males had either blunted or missing maxillary canines. While some canines were lost postmortem and remained with the skull, most were lost antemortem, as indicated by resorbed tooth sockets. Functionally, maxillary canines restrict the lateral movement of the opposing mandible, providing a sharpening mechanism to maintain chewing functionality and efficiency (Bramblett, 1969). When maxillary canines are removed, this restriction is diminished or eliminated, allowing greater lateral movement of the mandible during chewing, redistributing occlusal forces, and accelerating molar wear.

Biomechanical differences in musculoskeletal design may also help explain the observed wear differences. Male baboons have significantly larger maximum jaw gapes relative to jaw length compared to females, 110% versus 90%, respectively, resulting in decreased mechanical efficiency during chewing (Hylander, 2013). Recent comparative work across cercopithecoid primate species further showed that males, but not females, displayed strong and consistent correlations between canine height, jaw gape, and musculoskeletal traits that facilitate wide-mouth opening (Taylor et al., 2024). In fact, only males showed size-correlated decreases in jaw muscle leverage, which favors increased gape, but compromises bite force. These findings suggest that in males, selection favored morphological configurations that support canine display and intrasexual competition, with less emphasis on chewing performance (Taylor et al., 2024). As a result, males may need to perform more chewing cycles to achieve similar levels of oral processing as females, leading to greater cumulative wear, even under controlled dietary conditions. If display behaviors involving canine height and gape are key targets of selection in male cercopithecoids, these adaptations may carry functional costs to the dentition over time.

Taken together, these anatomical and behavioral insights indicate that multiple, overlapping factors likely contribute to the greater wear observed in male baboons. The most plausible explanations involve sexually dimorphic enamel thickness, canine removal, altered muscle architecture, and decreased bite force efficiency in favor of wide gape. Importantly, if gape- and display-related morphologies have shaped the evolution of the male cercopithecoid masticatory system, then caution is warranted when using these taxa as models for reconstructing feeding behavior in fossil hominins, as their craniofacial form may reflect social signaling demands more than dietary or ecological function (Taylor et al., 2024). Behavioral influences such as bruxism or food-softening practices may contribute, though direct observational data would be required to assess their influence. Further research using larger samples, behavioral monitoring, and enamel histology could help clarify the relative importance of these factors in shaping sex differences in dental wear.

### Wear progression is nonlinear and differs by sex

Mixed-effects regression models incorporating linear, quadratic, and cubic age terms (age, age^2^, and age^3^) indicated that wear progression follows a nonlinear trajectory, with wear decelerating later in life (Tables 7a and 7b). This pattern was more pronounced in males, whose Krueger-Scott scores and DER values began to diverge from those of females at approximately 150 months of age (Figure 4). These findings suggest that wear does not accumulate uniformly across the lifespan, but instead follows a phasic pattern, characterized by an initial phase of rapid wear, followed by a period of slower accumulation. This deceleration may reflect the progressive loss of enamel and increased exposure of softer, underlying dentin.

This trajectory was most clearly captured in Krueger-Scott scores. Isotonic regression plots showed increases in summed scores for both sexes during the first 100 months of life (Figure 4), while DER values remained relatively flat during this period and began to only rise around 150 months, coinciding with the point of male-female divergence (Figure 4). This pattern is consistent with findings from Phillips-Conroy et al. (2001), who observed in wild baboons that dentin exposure increased slowly after eruption, accelerated with continued wear, and decelerated once exposure was extensive. These results highlight a key advantage of the Krueger-Scott method: by capturing both enamel loss and dentin exposure, it detects early wear stages that DER alone may miss.

Quadrant-level analysis further illustrates this trend. Buccal cusps show the steepest increase in wear during early life, driving the overall pattern observed in summed scores. Although lingual cusps also show wear increases, their progression is more gradual, consistent with biomechanical differences in occlusal loading during early mastication (Figure 3).

These findings challenge the assumption, often implicit in cross-sectional studies, that wear progresses linearly with age (e.g., Bramblett, 1969; King et al., 2005; Wright et al., 2008). By reconstructing wear across three timepoints (inferred eruption, casting, and death), we show that wear trajectories are influenced by more than just elapsed time. They are shaped by changes in occlusal function, morphology, and potentially tertiary dentin responses. This approach highlights the unique value of longitudinal datasets, especially when sex-specific differences in wear timing and pattern are taken into account.

### Value of the Krueger-Scott method and implications for future research

Scott scores have been a foundational tool in dental anthropology for decades (Scott, 1979), and remain the standard for molar wear data collection in archaeological and paleoanthropological contexts (Buikstra and Ubelaker, 1994). Although various modifications have been developed for use in non-human primates (e.g. Avià et al., 2022; Conrad, 2023); we retained the original 1-10 ordinal scale and adapted the descriptions to reflect the occlusal morphology of cercopithecoid molars (Figure 1). Additionally, we incorporated the quadrant orientation and scoring order proposed by Shykoluk and Lovell (2010), enabling consistent application across individuals. These adjustments, together forming the Krueger-Scott method, proved effective for capturing regional wear patterns in our large baboon sample, and the standard quadrant scoring enabled more refined, functionally meaningful analyses of intra-tooth variation.

This modified approach offers several advantages for future studies. First, it provides a standardized yet anatomically sensitive method for collecting macrowear data in non-human primates using a scoring system that is already familiar to many researchers. As it aligns with recommended standards (Buikstra and Ubelaker, 1994), transitioning to this approach for cercopithecoid samples should be relatively seamless and enhance cross-study comparisons. Second, the method is efficient and accessible. Scoring the four quadrants of all specimens in the study required approximately two weeks, carried out by a single observer using only modified descriptions and no specialized equipment. This makes it especially well-suited for researchers operating with limited time, funding, or access to advanced imaging technology.

In contrast, collecting DER values for the full occlusal surface was a time-intensive process that took several months to collect on our large sample; approximately 100 hours of person-time were invested in collecting these data. Extending DER collection to each quadrant would be prohibitively laborious and impractical at scale and would still fail to capture early enamel wear. Therefore, we did not pursue this. We find that despite the more technologically advanced characteristics of DER values, the Krueger-Scott scoring method provides more nuanced insight to crown wear in addition to being a more efficient use of scientific resources. This insight is similar to an earlier investigation of tooth crown area for this same captive baboon population.

Hlusko et al. (2002) compared occlusal two-dimensional areas to the 2D area calculated as the product of mesiodistal length and buccolingual breadth. The residual heritability estimates were the same for both methods of crown area calculation. However, similar to what we found for DER versus Krueger-Scott scores, linear dimensions provide a more nuanced analysis despite being less technologically complicated (Hlusko et al., 2002; Hlusko et al., 2016).

By adapting an existing anthropological standard to suit non-human taxa, the Krueger-Scott method provides a replicable, morphologically informed tool for assessing wear across species and time periods, applicable to both extinct and extant cercopithecoids due to their shared bilophodont molar morphology (Avià et al., 2022). Its utility may also extend to other mammals with similar bilophodont dental architecture, such as lagomorphs and certain rodents. This approach offers both spatial resolution and quantitative depth, which only enhances our ability to interpret functional, developmental, and ecological aspects of dental wear in primates and beyond.

## Conclusion

This study demonstrates that dental wear progression in captive baboons, measured using a modified quadrant-level Scott scoring system and DER, follows consistent and interpretable patterns even under uniform dietary and environmental conditions. By comparing a traditional DER metric with a novel quadrant-level modification of Scott’s ordinal wear scoring system, the Krueger-Scott method, we evaluated not just the extent, but the regional dynamics of wear across the occlusal surface.

Our results show that wear accumulates in a non-uniform and functionally structured way, with buccal cusps wearing earlier and more quickly, and lingual cusps showing delayed but accelerated wear later in life. The Krueger-Scott method was especially effective in detecting these intra-occlusal dynamics, and revealed early enamel loss not detected by DER. In contrast, DER provided a later-stage summary of wear and required considerably more time to collect. Quadrant-level ordinal scoring thus offers a more efficient, morphologically informative, and accessible tool for assessing wear, particularly in bilophodont taxa.

We also found clear sex differences in wear progression, with males showing greater and faster wear despite their younger average age. This pattern likely reflects a combination of enamel structural variation, skeletal and biomechanical dimorphism, and management-related factors unique to captive populations. Together, these findings provide a new comparative baseline for interpreting molar wear and demonstrate the value of quadrant-level ordinal wear methods for identifying biologically meaningful variation, especially in studies of wild or fossil primates.

## Acknowledgements

This project was funded by the European Research Council within the European Union’s Horizon Europe (ERC-2021-ADG, Tied2Teeth) project number101054659. KLK acknowledges the LUC-INSPIRE microgrant program for travel funds to CENIEH and thanks Dr. Myra Laird for interesting conversations about baboon jaw biomechanics that informed the discussion. We thank Dr. James Cheverud for access to the baboon collection at Loyola University Chicago and Madeline Gustaf for help with intraoral scanning. The dental molds and casts were made for research supported by the National Science Foundation Division of Behavioral and Cognitive Sciences Grants 0500179, 0616308, and 0130277. This investigation used resources that were supported by the Southwest National Primate Research Center grant P51 OD011133 from the Office of Research Infrastructure Programs, National Institutes of Health. Many thanks to the Southwest National Primate Research Center for access and support throughout this long-term study of baboon dental variation. We thank all of the non-author members of the Tied2Teeth project for project support.

## Author Contribution Statement (CRediT)

Kristin Krueger

Conceptualization-Equal, Formal analysis-Equal, Investigation-Equal, Methodology-Lead, Validation-Supporting, Visualization-Supporting, Writing - original draft-Lead, Writing - review & editing-Lead

Ian Towle

Conceptualization-Equal, Formal analysis-Supporting, Investigation-Equal, Methodology-Supporting, Resources-Equal, Visualization-Supporting, Writing - original draft-Equal, Writing - review & editing-Supporting

Gregory Matthews

Formal analysis-Lead, Methodology-Supporting, Software-Lead, Visualization-Lead, Writing - review & editing-Equal

Ana Álvarez Fernández

Data curation-Supporting, Investigation-Supporting, Methodology-Supporting

Leslea Hlusko

Conceptualization-Equal, Data curation-Equal, Formal analysis-Supporting, Funding acquisition-Lead, Methodology-Supporting, Project administration-Lead, Resources-Lead, Supervision-Lead, Writing - review & editing-Supporting

## Data Availability Statement

The data that support the findings of this study are openly available in Dryad at the Reviewer URL: http://datadryad.org/share/kacvtLc_KnGSRnHCqXIv56Ew2dvodrXCQYD4TNVXZJ4

