## Supplementary material for "Tracking molar wear in captive baboons: sex and age effects using a modified Scott scoring system": Tables

**Table 1.** General information about the study sample, including the number of individuals, average age at the cast wear score and death wear score, and average time between recordings, split by sex (all, female, and male). All ages and time are in months.

| Sample | Value |
| --- | --- |
| Number of individuals | 201 |
| Number of males | 51 |
| Number of females | 150 |
| Average age at cast wear score (all) | 224.0 |
| Average age at cast wear score (female) | 233.5 |
| Average age at cast wear score (male) | 196.1 |
| Average age at death wear score (all) | 253.2 |
| Average age at death wear score (female) | 263.0 |
| Average age at death wear score (male) | 224.4 |
| Average time between recordings (all) | 29.1 |
| Average time between recordings (female) | 29.4 |
| Average time between recordings (male) | 28.3 |

**Table 2a.** Mean and standard deviation of the cast and death wear Krueger-Scott scores for each tooth position (mesiobuccal, distobuccal, mesiolingual, distolingual) and all quadrants summed. This includes overall averages as well as averages by sex.

| Quadrant | Cast Scott score  mean | Cast Scott score  std | Death Scott score  mean | Death Scott score  std |
| --- | --- | --- | --- | --- |
| **All individuals** |  |  |  |  |
| Summed score | 18.9 | 4.92 | 22.6 | 4.82 |
| Distobuccal | 5.31 | 1.15 | 6.06 | 1.09 |
| Mesiobuccal | 5.85 | 0.98 | 6.50 | 0.96 |
| Distolingual | 3.62 | 1.87 | 4.69 | 1.80 |
| Mesiolingual | 4.09 | 1.79 | 5.35 | 1.73 |
| **Females** |  |  |  |  |
| Summed score | 18.0 | 4.70 | 21.6 | 4.35 |
| Distobuccal | 5.21 | 1.15 | 5.93 | 1.03 |
| Mesiobuccal | 5.81 | 0.99 | 6.41 | 0.88 |
| Distolingual | 3.19 | 1.69 | 4.24 | 1.58 |
| Mesiolingual | 3.79 | 1.75 | 5.01 | 1.57 |
| **Males** |  |  |  |  |
| Summed score | 21.5 | 4.70 | 25.5 | 4.98 |
| Distobuccal | 5.61 | 1.11 | 6.43 | 1.17 |
| Mesiobuccal | 5.98 | 0.95 | 6.78 | 1.10 |
| Distolingual | 4.88 | 1.80 | 6.00 | 1.78 |
| Mesiolingual | 4.98 | 1.63 | 6.33 | 1.82 |

**Table 2b.** Mean and standard deviation of the cast and death wear DER. This includes overall averages as well as averages by sex.

| Overall | Cast DER  mean | Cast DER score  std | Death DER score  mean | Death DER score  std |
| --- | --- | --- | --- | --- |
| **All individuals** | 0.118 | 0.107 | 0.151 | 0.126 |
| **Females** | 0.106 | 0.09 | 0.132 | 0.104 |
| **Males** | 0.152 | 0.133 | 0.206 | 0.163 |

**Table 3a.** Two-sample dependent t-tests for hypothesis 1, Krueger-Scott scores. Bolded value denotes a statistically significant *p* value.

| **Quadrant** | **t** | ***df*** | ***p*** |
| --- | --- | --- | --- |
| Distobuccal skeletal- Distobuccal cast | -15.408 | 200 | **<0.0001** |
| Mesiobuccal skeletal-Mesiobuccal cast | -13.803 | 200 | **<0.0001** |
| Distolingual skeletal-Distolingual cast | -14.071 | 200 | **<0.0001** |
| Mesiolingual skeletal-Mesiolingual cast | -15.343 | 200 | **<0.0001** |

**Table 3b**. ANOVA for hypothesis 1, DER. Bolded value denotes a statistically significant *p* value.

| **DER** | **t** | **estimate** | ***p*** |
| --- | --- | --- | --- |
| DER skeletal-DER cast | 5.877 | .0331 | **<0.0075** |

**Table 4.** Spearman correlation coefficients for hypothesis 2. The Bonferroni corrected cut-off for significance was 0.0017857. All *p-*values were well below this value.

|  | DL cast | DL  skeletal | ML cast | ML  skeletal | MB  skeletal | MB cast | DB  skeletal | DB cast |
| --- | --- | --- | --- | --- | --- | --- | --- | --- |
| DL cast | **1** | **0.84** | **0.74** | **0.70** | **0.49** | **0.49** | **0.49** | **0.50** |
| DL skeletal | **0.84** | **1** | **0.66** | **0.74** | **0.55** | **0.49** | **0.61** | **0.53** |
| ML cast | **0.74** | **0.66** | **1** | **0.82** | **0.51** | **0.61** | **0.49** | **0.53** |
| ML skeletal | **0.70** | **0.74** | **0.82** | **1** | **0.62** | **0.59** | **0.56** | **0.52** |
| MB skeletal | **0.49** | **0.55** | **0.51** | **0.62** | **1** | **0.76** | **0.72** | **0.67** |
| MB cast | **0.49** | **0.49** | **0.61** | **0.59** | **0.76** | **1** | **0.71** | **0.78** |
| DB skeletal | **0.49** | **0.61** | **0.49** | **0.56** | **0.72** | **0.71** | **1** | **0.80** |
| DB cast | **0.50** | **0.53** | **0.53** | **0.52** | **0.67** | **0.78** | **0.80** | **1** |

**Table 5a**. Hotelling’s T^2^ tests for hypothesis 3, bolded value denotes statistically significant *p* value.

|  | Hotelling’s T^2^ | *p*-value |
| --- | --- | --- |
| Cast by sex | 42.938 | **<0.0001** |
| Skeletal by sex | 46.791 | **<0.0001** |

**Table 5b.** Welch’s t-tests for hypothesis 3. The Bonferroni corrected cut-off for significance was 0.00625. Bolded value denotes a statistically significant *p* value.

|  | t | df | *p*-value | Mean in females | Mean in males |
| --- | --- | --- | --- | --- | --- |
| MB cast by sex | -1.118 | 89.591 | 0.2665 | 5.807 | 5.980 |
| DB cast by sex | -2.203 | 88.72 | 0.0302 | 5.207 | 5.608 |
| ML cast by sex | -4.430 | 92.159 | **<0.0001** | 3.787 | 4.980 |
| DL cast by sex | -5.885 | 82.279 | **<0.0001** | 3.193 | 4.882 |
| MB skeletal by sex | -2.219 | 73.084 | 0.0296 | 6.411 | 6.784 |
| DB skeletal by sex | -2.701 | 78.207 | 0.0085 | 5.933 | 6.431 |
| ML skeletal by sex | -4.629 | 76.952 | **<0.0001** | 5.013 | 6.333 |
| DL skeletal by sex | -6.279 | 78.517 | **<0.0001** | 4.24 | 6.000 |

**Table 6**. DER at cast and at death by sex. Mixed effects regression model.

|  | *p*-value | Mean in females | Mean in males |
| --- | --- | --- | --- |
| DER cast by sex | **0.00685** | 5.807 | 5.980 |
| DER skeletal by sex | **0.00228** | 5.207 | 5.608 |

**Table 7a.** Mixed-effects models *p*-values and Intraclass correlation (ICC) results for hypothesis 4, Krueger-Scott scores. Bolded value denotes a statistically significant *p* value.

| **Quadrant** | ***p*-value** | **ICC** |
| --- | --- | --- |
| Mesiobuccal | **<0.0001** | 0.3120 |
| Distobuccal | **<0.0001** | 0.3303 |
| Mesiolingual | **<0.0001** | 0.2979 |
| Distolingual | **<0.0001** | 0.3382 |
| Summed score | **<0.0001** | 0.3322 |

**Table 7b**. Mixed-effects models *p*-values and Intraclass correlation (ICC) results for hypothesis 4, DER. Bolded value denotes a statistically significant *p* value.

| **DER** | ***p*-value** | **ICC** |
| --- | --- | --- |
| DER by observer | **<0.0001** | 0.0023 |
| DER by baboon ID | **<0.0001** | 0.5461 |
